# The principles of expected and realised genetic relatedness among individual honeybees

**DOI:** 10.1101/2024.05.26.595938

**Authors:** Laura Strachan, Jernej Bubnič, Gertje Petersen, Gregor Gorjanc, Jana Obsteter

## Abstract

Monitoring honeybee genetic variability is essential to manage global and local genetic diversity. Coefficients of relatedness are regularly used to measure genetic similarity within and between populations and their individuals. Although the haplo-diploid inheritance of honeybees is well understood, interpreting the various types of relatedness coefficients based on pedigree and genotype data is a challenge for researchers and practitioners in honeybee breeding. To demonstrate the principles of genetic relatedness in honeybees and its different individual-based estimators, we simulated three honeybee populations each containing 400 colonies over 10 years using the stochastic simulator SIMplyBee. We kept two populations closed and hybridised the third one by importing drones from one of the closed populations. We evaluated the relatedness between individuals within a colony, between queens of the same population, and between queens of different populations. We calculated three types of relatedness: expected identity by descent using pedigree information, realised identity by descent using pedigree and genotype information, and identity by state using genotype information. Our results demonstrated an alignment of mean relatedness across different types when calculated using the same founder population, regardless of their data source. Identity by state relatedness varied significantly when calculated with different founder populations. Although this is an expected result, it shows that caution is needed when comparing values between studies using different populations with different allele frequencies. We expectedly showed increased relatedness over time in closed populations and decreased in the hybrid population. Our results underscore the significance of understanding the methodology for computing relatedness coefficients.

## 1 Introduction

Monitoring and managing genetic variability is crucial to safeguarding genetic diversity in local and global populations. The loss of genetic variability diminishes the fitness of extant populations and their long-term adaptability, which are essential in dynamic environments. Furthermore, conscious management of genetic variability is central to breeding programmes to facilitate short-term genetic improvements aligned with current breeding goals, as well as long-term genetic improvements in anticipation of future breeding goals.

Conservation by utilisation is considered the most effective approach for supporting and managing genetic variability. A crucial aspect of this strategy involves breeding programmes to enhance economically important traits while managing diversity, as intensive selection can lead to its decline (Falconer and Mackay, 1996; Charlesworth and Charlesworth, 1999; Uzunov *et al*., 2017). Inbreeding depression, characterised by reduced individual genetic variability, can result in reduced performance. This is attributed to factors such as the expression of partially recessive deleterious mutations or increased homozygosity at loci expressing overdominance (Falconer and Mackay, 1996; Charlesworth and Charlesworth, 1999). Furthermore, increased relatedness and inbreeding enhance genetic drift, driving random changes in allele frequency and linkage, potentially resulting in unforeseen changes in economically important traits. To manage such losses of genetic diversity, it is essential to optimise parental selection and mating allocations based on relatedness between individuals (Wright, 1921).

The Western Honeybee (*Apis mellifera* L.) is a globally important economic species that plays a vital role in pollination services and food production (Klein *et al*., 2007; Gallai *et al*., 2009). Recent research has sparked a growing interest in monitoring and managing genetic diversity within honeybee populations, due to emerging studies demonstrating a decline in genetic variability of honeybee subspecies when comparing historical and current samples (Espregueira Themudo *et al*., 2020). The genetic variability of honeybees is impacted by abiotic factors, including climate changes, the reduction of suitable habitats, agricultural intensification with increased agrochemical usage, and an increasing burden of pathogens and parasites (Zayed, 2009; Jaffé *et al*., 2010).

Several biotic factors also affect the genetic diversity of honeybee populations, including haplo-diploidy, the complementary sex determiner (*CSD*) locus, the importation of queens, and selective breeding practices. Honeybees have a unique haplo-diploid sex-determining system, whereby diploid females stem from fertilized eggs and haploid males arise from unfertilized eggs, with the *CSD* locus regulating sex differentiation (Beye *et al*., 2006). Fertilised eggs heterozygous at the *CSD* locus develop into diploid females, whereas homozygotes produce sterile or non-viable diploid drones (Woyke, 1962). These mechanisms effectively prevent inbreeding, thus maintaining genetic diversity in honeybee populations. An additional mechanism to prevent inbreeding is polyandry, whereby a virgin queen mates with multiple males during her mating flights. Consequently, three types of sisters can be present in the diploid offspring (half-sisters, full-sisters, and super-sisters) dependent on the relatedness between fathers’ sperm used for egg fertilisation. Half-sisters have unrelated fathers, full-sister have related fathers (brother drones produced by the same queen), and super-sisters share the same father (Crow and Roberts, 1950; Crozier, 1970; Laidlaw Jr and Page Jr, 1984). Introgressive hybridisation, driven by natural mating at geographic subspecies borders or import of nonnative subspecies, increases genetic diversity as well by mixing different populations, although this can lead to a decline of a native population (De la Rúa *et al*., 2013; Espregueira Themudo *et al*., 2020). In contrast, selective breeding practices generally decrease genetic variation by rearing preferred families, though such practices can also increase the frequency of rare alleles and transitionally increase genetic variance (Crow, 2010). Furthermore, most of genetic variance reduced due to selection within a generation is restored via recombination between generations (Bulmer, 1971; Lara *et al*., 2022).

Reduced genetic variability negatively affects honeybee population by increasing brood losses (Woyke, 1962; Zayed *et al*., 2006), compromising colony efficiency (Oxley and Oldroyd, 2010; Espregueira Themudo *et al*., 2020), leading to inbreeding depression for fitness traits (Antolin, 1999; Henter, 2003; Zayed, 2009), and impairing long-term adaptive capacity, in turn increasing the risk of extinction.

To understand genetic diversity within a population, Wright (1921) introduced coefficients of relationship and inbreeding within a known pedigree. He based this work on diploid individuals and additive gene action such that the genetic value of an individual *i* (*a_i_*) is the sum of values of its two genomes: *a_i_* = *a_i,_*_1_ + *a_i,_*_2_. Wright’s coefficient of relationship is the correlation between the genetic values of two individuals, denoted as *R_i,j_* = *Cor* (*a_i_, a_j_*), while Wright’s coefficient of inbreeding is the correlation between the genetic values of the two genomes of an individual, expressed as *F_i_* = *Cor* (*a_i,_*_1_*, a_i,_*_2_). Wright (1921) showed how to calculate these coefficients by tracing all pedigree lineages between relatives. Cotterman (1940) and Malécot (1948) have later shown that Wright’s coefficients measure the expected identity by descent (IBD) of individual’s genomes within a pedigree relative to the pedigree founders, while observing individuals’ genomes would enable measuring identity by state (IBS).

Emik and Terrill (1949) developed a recursive algorithm to calculate the relatedness coefficients for all pairs of individuals within a pedigree. This algorithm works with an *n × n* covariance coefficient matrix **A** for *n* individuals and sets this matrix to an identity matrix between pedigree founders. Diagonal elements of the matrix are defined as: 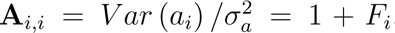, with values in the range of [1, 2] since *F_i_* is the correlation coefficient. Off-diagonal elements of the matrix are defined as: 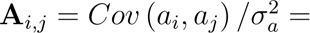^1^*/*_2_ (*Cor* (*a_i,_*_1_*, a_j,_*_1_) + *Cor* (*a_i,_*_1_*, a_j,_*_2_) + *Cor* (*a_i,_*_2_*, a_j,_*_1_) + *Cor* (*a_i,_*_2_*, a_j,_*_2_)), with values in the range of [0, 2]. *F_i_* is estimated as half the off-diagonal element between the parents of *i*: ^1^*/*_2_**A***_m_*_(_*_i_*_)_*_,f_*_(_*_i_*_)_. Off-diagonal elements are estimated as: **A***_i,j_* = ^1^*/*_2_ **A***_i,m_*_(_*_j_*_)_ + **A***_i,f_*_(_*_j_*_)_. Completing the matrix **A** enables calculating Wright’s inbreeding coefficients *F_i_* and relationship coefficients *R_i,j_*. However, it is now common to use just elements of **A**, which Henderson (1976) named as the numerator relationship matrix.

Pedigree-based relatedness coefficients exhibit two important limitations. First, they measure expected relatedness between individuals, wherein the expectation is predicted based on the recombination and segregation processes of genomes between parents and progeny. However, there is substantial variation in the realised relationships between pairs of relatives due to these stochastic processes (Hill and Weir, 2011). Second, pedigree-based coefficients measure relatedness relative to the assumed unrelated founder population at the start of the pedigree. Hence pedigree-based coefficients are limited by the unobserved inheritance of genomes, assumed unrelated founders, and pedigree depth. Nevertheless, pedigree-based relatedness measures expected IBD, which is expectation of the realised IBD, which is in turn expectation of the IBS, all relative to the base population of pedigree founders.

With the availability of genome-wide genotype (genomic) data, we can nowadays observe actual relatedness between individuals due to IBS, which is ultimately driven by mutations and their recombination and segregation across generations (Visscher *et al*., 2006; VanRaden, 2008; Hill and Weir, 2011). Denote **w***_i_* as a row-vector of genotypes of individual *i* at *n_m_* markers encoded as the number of alternative alleles. Similarly to the pedigree-based numerator relationship matrix, elements of genotype-based numerator relationship matrix are calculated from the observed covariance between genotypes of pairs of individuals: **w***_i_***w***^T^* (VanRaden, 2008). In this calculation, the genotypes are often centred and scaled to present genotype-based relatedness relative to a chosen base/reference population (VanRaden, 2008; Yang *et al*., 2010; Speed and Balding, 2015). The centring is based on allele frequency calculated from the chosen base/reference population. Coupled with pedigrees, the availability of genomic data also allows observing the realised IBD, by reconstructing the inheritance of genomes between generations of a pedigree (Elston and Stewart, 2008; Whalen *et al*., 2018).

Estimating relatedness among honeybees has to take into account their haplo-diploid inheritance. Grossman and Eisen (1989) and Grossman and Fernando (1989) developed a general method for calculating pedigree-based relatedness for the X chromosome, which has haplo-diploid inheritance and can also be used for individual honeybees. Recently, Druet and Legarra (2020) developed a general method for calculating genotype-based relatedness matrix for the X chromosome. They also presented a method for calculating a “hybrid” relatedness matrix that combines pedigree and genotype data when some individuals are not genotyped.

Although the genome-wide genotype data is now becoming widely available for honey-bees, the current landscape of honeybee genetic studies reveals a gap in the utilisation and comparison of methodologies in constructing genomic relationship matrices (Tarpy *et al*., 2023). While some GWAS studies briefly mention the calculation of genotype-based re-latedness, they lack explicit discussion on the methodologies employed and leave room for assumptions (Avalos *et al*., 2020; Eynard *et al*., 2024). In the majority of these studies, the allele frequency used are derived from pooled samples typically collected from a wide range of locations, possibly encompassing different subspecies and their hybrids. Consequently, this large genetic diversity challenges the comparison of the resulting genotype-based relatedness to classic pedigree-based relatedness. Therefore, there is a need for studies to explicitly de-lineate the methods used in calculating relatedness among individual honeybees, particularly focusing on allele frequency and IBS calculations, to estimate the impact of chosen reference populations on the relatedness values. This clarification will aid in better study comparisons and improve the management of genetic variability in honeybee populations.

The overarching aim of this contribution is to demonstrate genetic relatedness among individual honeybees within a colony, within a population, and comparatively between pop-ulations, using simulation. Specifically, we aim to: (i) demonstrate the principles of genetic inheritance in honeybees by evaluating relatedness within a colony; (ii) compare relatedness calculated from different sources of information and using different methodology; and (iii) measure the impact of hybridisation between honeybee populations.

## 2 Materials and Methods

We used the stochastic simulator SIMplyBee Obšteter *et al*. (2023) to simulate a 10-year cycle of two closed honeybee populations and their hybrid. We inspected whole-genome and *CSD* locus relatedness values within a colony, between queens of the same population, and between queens of different populations. We computed three types of relatedness values based on: (i) expected identity by descent (eIBD) information, (ii) realized identity by descent (rIBD) information, and (iii) identity by state (IBS) information.

### 2.1 Simulation of the honeybee population

#### 2.1.1 Simulation of founder population

We simulated the founder haplotypes using the Markovian Coalescent Simulator (MaCS; (Chen *et al*., 2009)) that is integrated into SIMplyBee via AlphaSimR (Gaynor *et al*., 2021). We simulated 400 founder whole-chromosome haplotypes of the “Carniolan honeybee” (*Apis mellifera carnica*) and 800 founder whole-chromosome haplotypes of the “European dark bee” (*Apis mellifera mellifera*), according to the demographic model described in Wallberg et al. (2014) (Wallberg *et al*., 2014). The mutation rate was set to 3.4e-09 (Yang *et al*., 2015), and the recombination rate to 2.3e-07 (Beye *et al*., 2006). We simulated 16 chromosomes and retained 1000 segregating sites per chromosome to monitor the relatedness. In addition, we simulated the *CSD* locus as a non-recombining region of DNA, located on the third chromosome, with 128 possible alleles allocated at random (Zareba *et al*., 2017; Obšteter *et al*., 2023). From the founder haplotypes, we created a founder population of unrelated diploid virgin queens, 400 *A.m.carnica* and 800 *A.m.mellifera*. We subsequently edited the *CSD* locus for all the virgin queens to be heterozygous and hence viable.

Throughout the remainder of the simulation we have removed any diploid individuals homozygous at the *CSD* locus. The founder queens were assigned to three populations, each with 400 virgin queens: two *A.m.mellifera* populations, one intended for within-species mating (Mel) and one for hybridisation (MelCross), and one *A.m.carnica* population (Car) for within-subspecies mating (Figure 1). Within each population, we randomly selected 300 virgin queens as the virgin queens of future colonies, and 100 as the drone-producing queens (DPQs). We initiated each population by mating the 300 virgin queens with an average of 15 drones produced by the 100 DPQs, and creating 300 colonies.

**Figure 1:**
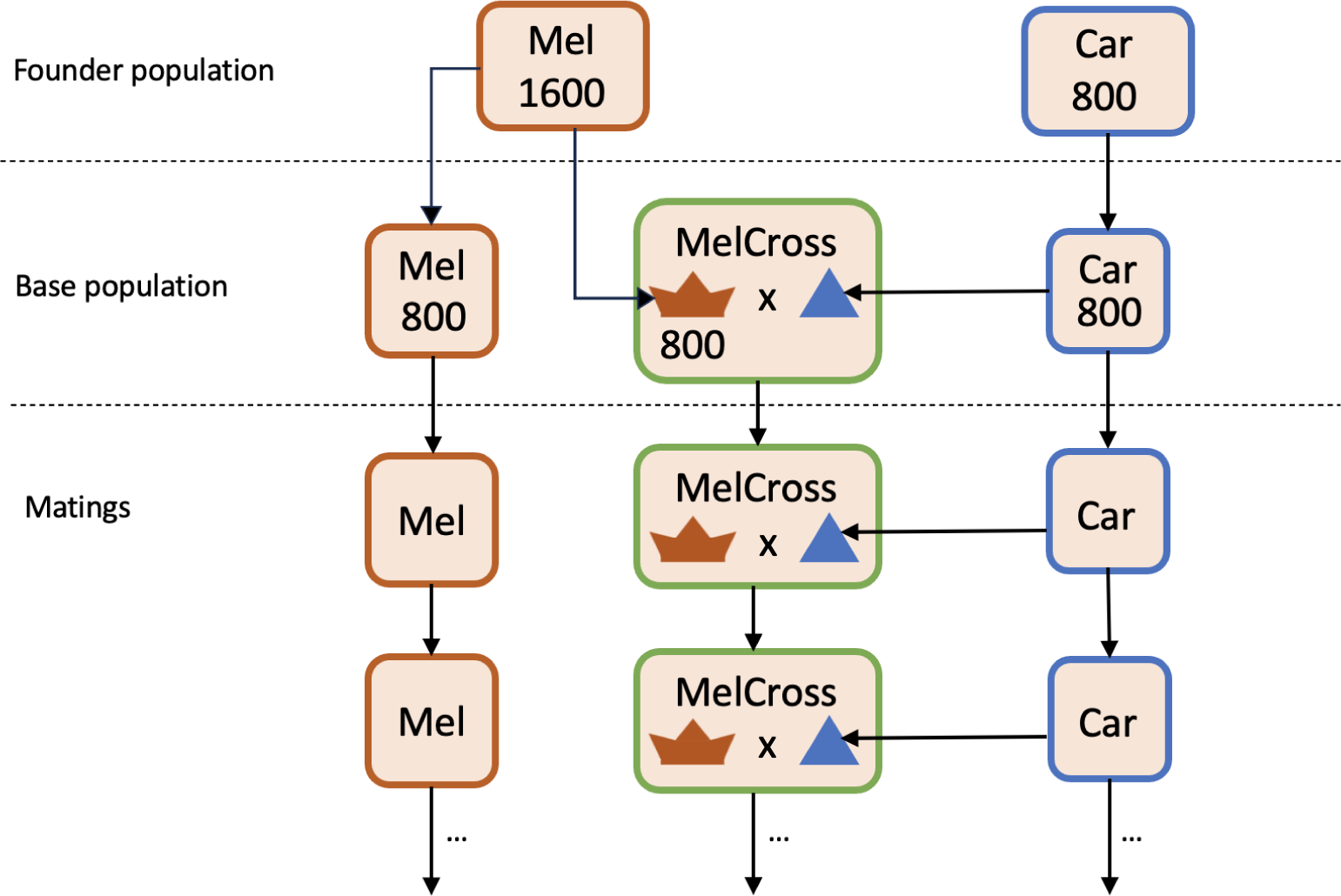
A flow diagram of the simulation, showing the founder population, base population and descending generations with the pathway of matings highlighted with arrows. 1600 *A.m.mellifera* and 800*A.m.carnica* founder haplotypes are used to create three founder populations of unrelated virgin queens, 400 *A.m.carnica* and 400 *A.m.mellifera* and a hybrid MelCross population contain 400 *A.m.mellifera* virgin queens. The *A.m.mellifera* and *A.m.carnica* populations had a closed population breeding system, while the hybrid MelCross population virgin queens were mated with *A.m.carnica* drones.

#### 2.1.2 10-year breeding programme

We conducted a 10-year simulation, where one year represented the annual series of events of a honeybee colony with three periods that represented spring, summer, and winter. The simulation incorporated key colony events including split, swarm, supersedure, and collapse, with probabilities varying based on the specific period as outlined in Table 1. For details on how SIMplyBee simulates colony events, see Obšteter *et al*. (2023).

**Table 1:**
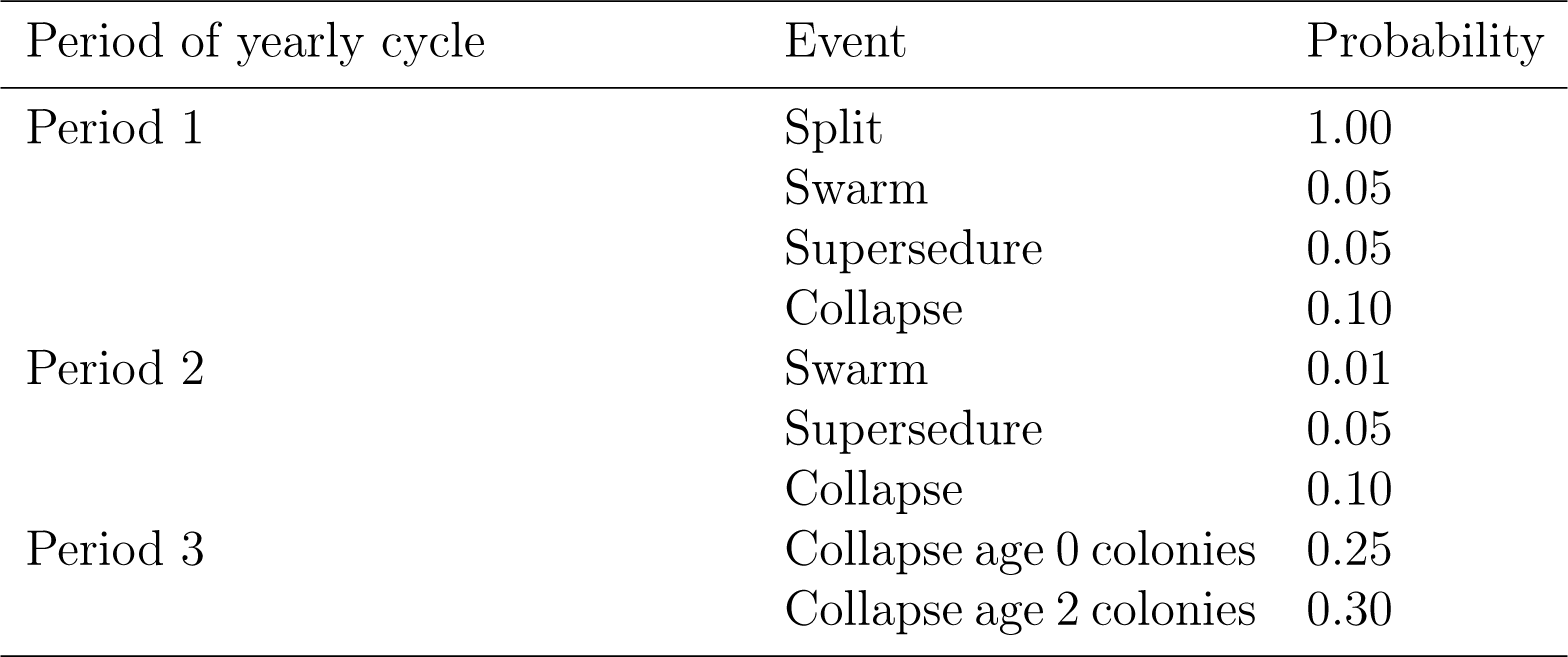
Probability of events during the yearly cycle.

| Period of yearly cycle | Event | Probability |
| --- | --- | --- |
| Period 1 | Split | 1.00 |
|  | Swarm | 0.05 |
|  | Supersedure | 0.05 |
|  | Collapse | 0.10 |
| Period 2 | Swarm | 0.01 |
|  | Supersedure | 0.05 |
|  | Collapse | 0.10 |
| Period 3 | Collapse age 0 colonies | 0.25 |
|  | Collapse age 2 colonies | 0.30 |

In period 1, all colonies were built up by adding 50 drones and 10 workers followed by a managed split. Subsequently, a subset of colonies with one or two-year-old queens experienced additional events based on probabilities (Table 1). The split and swarm events led to an increase in the number of simulated colonies, generating a “split” or “swarmed” colony with a new queen and a parent “remnant” colony with the previous queen. Given the managed nature of a split, the virgin queens of “split” colonies were replaced with donor virgin queens generated from 5 one-year-old queens, representing selection by beekeepers. All virgin queens resulting from split, swarm, and supersedure events were mated with drones from a drone congregation area (DCA) that consisted of drones from colonies with one-or two-year-old queens. Mating schemes differed between closed and hybrid populations, the former mating only with drones of the same population and the latter mating with a mix of 50% of their own MelCross drones and 50% of Car drones (Figure 1).

Period 2 varied only slightly from Period 1. In Period 2, no managed splits were performed and all colony events occured with probabilities outlined in Table 1. Virgin queen matings were identical to that of Period 1.

As Period 3 represents winter, the only colony event that took place was a collapse to mimic winter colony losses. The probability of a collapse was determined by the queens’ age (Table 1).

To maintain a population size of 300 colonies, we kept all colonies with one-year-old queens. If these summed up to less than 300, we “topped-up” populations with remnants from splits. Any excess colonies were removed from the population, which represented sale or export. Any colonies containing queens over the age of two were removed to simulate a more realistic managed honeybee queen life-cycle.

### 2.2 Calculations of relatedness

We used three relatedness metrics based on different sources of information: (i) expected IBD (eIBD) by using pedigree data; (ii) realised IBD (rIBD) by using pedigree and recombination data; and (iii) IBS by using genotype data. We express relatedness based on the pedigree or genomic numerator relationship matrix, which is equal to twice the coefficient of kinship (Wright, 1921; Henderson, 1976), specific for haplo-diploid inheritance (Grossman and Eisen, 1989). Hence, we report relatedness values in the range of [0, 2].

We calculated the eIBD relatedness based on the collected pedigree of all simulated individuals over all the populations and years. To manage the compute and memory requirements, we have first used R package *nadiv* (Wolak, 2012) to compute the sparse inverse of pedigree-based relationship matrix for haplo-diploid inheritance following Grossman and Fernando (1989). We next used the indirect method of Colleau (2002) to calculate the dense eIBD relationship matrix only for individuals of interest.

We calculated the rIBD relatedness based on the origin of alleles. Alleles’ origins were tracked with SIMplyBee/AlphaSimR since founders by labeling the alleles of founders’ whole-chromosome haplotypes with unique codes (from 1 to twice the number of founders) and tracking their recombination and segregation through the pedigree. We then calculated the rIBD relationship matrix for individuals of interest as twice the proportion of shared alleles IBD between pairs of individuals using the SIMplyBee function *calcBeeGRMIbd*.

We calculated the IBS relatedness from genotype data on 1000 segregating sites per chromosome. We used SIMplyBee function *calcBeeGRMIbs* that implements the method described by Druet and Legarra (2020) for computing haplo-diploid relationship matrix. To demonstrate the impact of different allele frequencies on the IBS relatedness, we used: (i) a colony’s own allele frequency (ColonyAF) estimated from all individuals within a colony, (ii) allele frequencies of subspecies-specific founder queens (SingleAF), (iii) allele frequencies of founder queens of both subspecies, *A.m.mellifera* and *A.m.carnica*, combined (MultiAF), or (iv) allele frequency of 0.5 (0.5AF).

We show all three relatedness metrics among individuals in year 1 and 10 and in three “dimensions”: (i) between individual honeybees of different castes within a colony, (ii) between queens of the same population, and (iii) between queens across different populations. For the within-colony “dimension”, we have first randomly selected a colony in each of the populations in years 1 and 10 and computed the within-colony relatedness between the queen, 1000 workers, 200 drones, any virgin queens present after colony events, and all father-drones that have mated with the queen. For the within-population “dimension”, we have computed the relatedness between all the queens of the same population (Car, Mel, and MelCross). We inspected the impact of hybridisation by inspecting separately the off-diagonal and diagonal elements of relationship matrices, which respectively represent the relatedness between same-population queens and the relatedness of the queens to themselves. For the between-population “dimension”, we have inspected the relatedness between queens of different pop-ulations by analysing the off-diagonal elements of relationship matrices.

R code for the simulations, relatedness calculations, and data analysis is available at https://github.com/HighlanderLab/lstrachan_honeybee_relatedness.

## 3 Results

### 3.1 Results Summary

Principles of honeybee genetic inheritance were demonstrated by inspecting relatedness within a colony in year 1 and 10 of closed population breeding.

Overall our results showed that the expected and realised IBD and IBS relatedness, over the 10-year simulation period, exhibit an increase in closed populations and a decrease in the hybrid population. Relatedness in all castes was shown to increase with years of closed population breeding due to increasing relatedness between queens and fathers. This closed population breeding increased the relatedness the most between workers and the least between drones. When comparing different metrics, calculations using the same pedigree founder population and reference population resulted in an alignment of mean relatedness, regardless of them being computed with pedigree or genotype data. However, when we assumed different reference populations for IBS relatedness we observed a large amount of variation in the resulting values, concluding that the results obtained can be easily misinter-preted depending on the allele frequency used in the calculations. These findings suggest that special attention has to be paid to understanding the used methodologies when interpreting the results.

### 3.2 Relatedness within a colony

We start by comparing relatedness between different caste individuals within a purebred colony, with relatedness based on eIBD, rIBD, and IBS calculated using allele frequencies of the founder *A.m.carnica* queens (SingleAF). We first present the results for year 1 by examining the worker-to-worker (WW) relatedness, followed by workers-to-drones (WD), and drones-to-drones (DD) relatedness. To inspect the effects of closed-population breeding, we next compare these results to year 10. We visualise these results in Figure 2 and present the corresponding mean and standard deviation of each relatedness metric for different types of sisters in Table 2.

**Figure 2:**
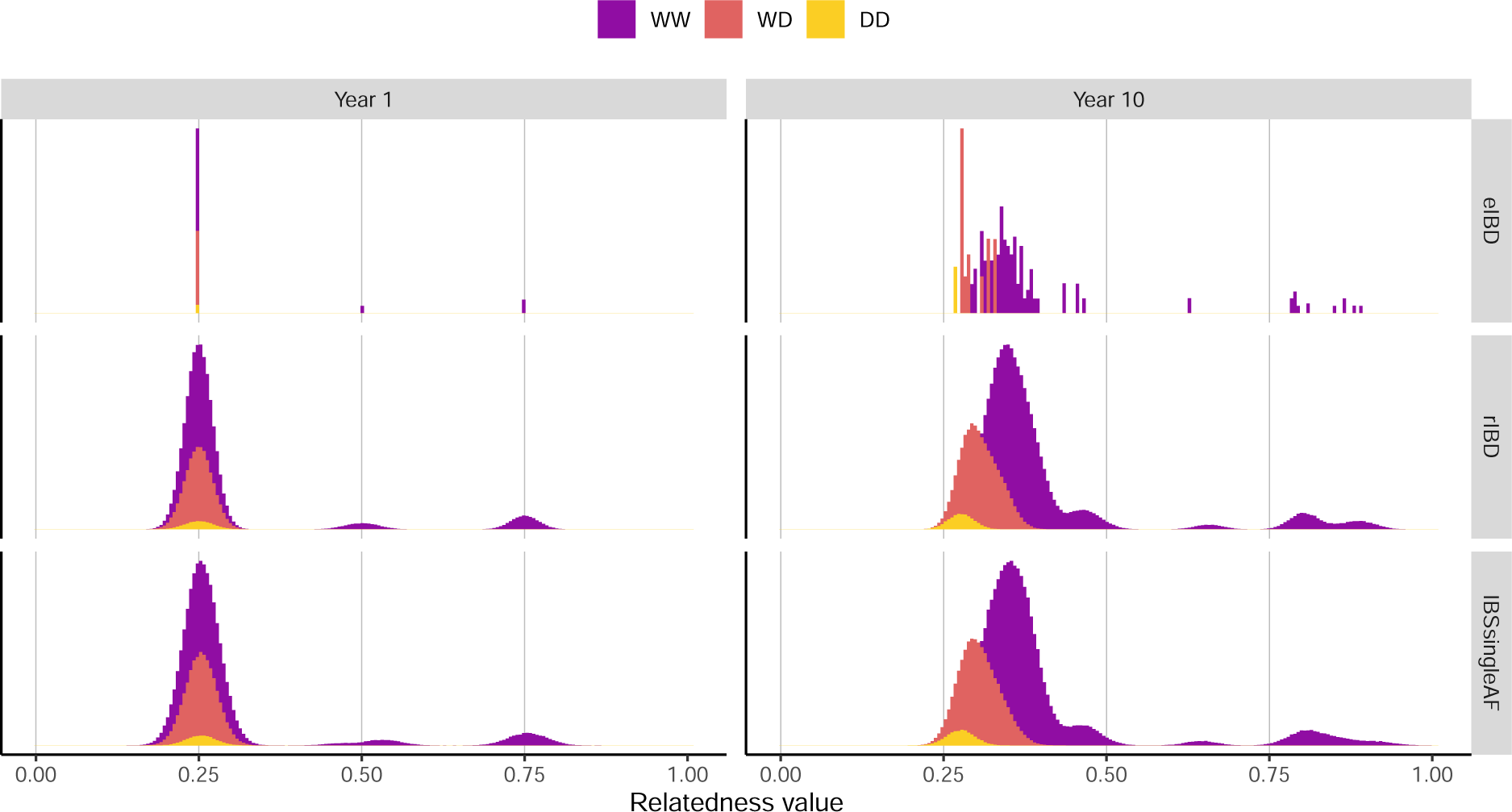
Expected and realised identity-by-descent (IBD) and identity-by-state (IBS) re-latedness within a *A.m.carnica* colony in year 1 (left) and 10 (right) of the simulation. IBS was calculated using allele frequencies of the founder *A.m.carnica* queens (SingleAF). The figure shows relatedness between workers-to-workers (WW), workers-to-drones (WD), and drones-to-drones (DD). Vertical lines at 0.00, 0.25, 0.50, and 0.75 represent the expected within-colony relatedness of a non-inbred colony to facilitate comparisons.

**Table 2:**
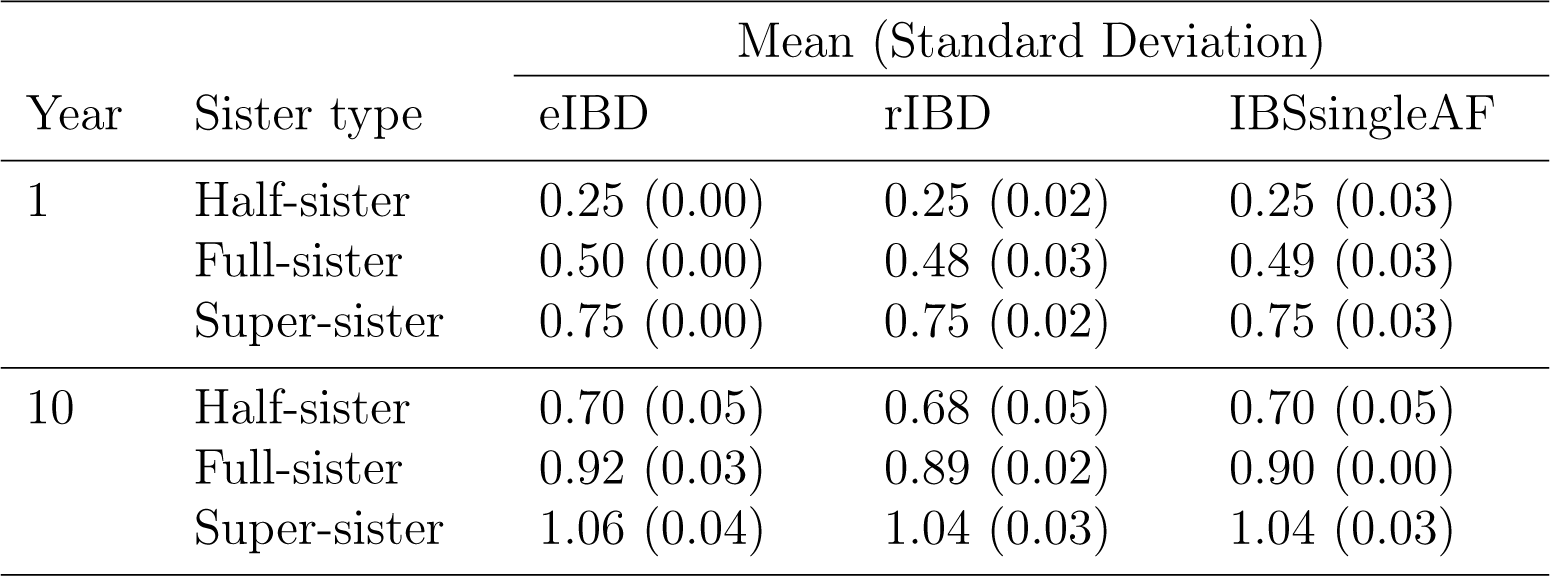
Mean and standard deviation of expected and realised identity-by-descent (IBD) relatedness and identity-by-state (IBS) relatedness between workers by sister type within a *A.m.carnica* colony in year 1 and year 10 of the simulation. IBS was calculated using allele frequencies of the founder *A.m.carnica* queens (SingleAF).

In year 1, eIBD relatedness between workers (WW) was seen at three distinct values (Figure 2). These values correspond to the different types of sisters in a colony, with 0.25 denoting half-sisters with unrelated fathers, 0.50 denoting full-sisters whose fathers are brothers (from the same queen), and 0.75 denoting super-sisters who share the same father. These values correspond to the literature on relatedness between honeybee workers (Crow and Roberts, 1950; Crozier, 1970; Laidlaw Jr and Page Jr, 1984) and more general literature on haplo-diploid inheritance (Grossman and Eisen, 1989; Druet and Legarra, 2020). eIBD relatedness between workers and drones (WD) within a colony is 0.25 (Figure 2) because they are only related through the maternal side. Lastly, eIBD relatedness between drones (DD) is 0.25 (Figure 2), again because all of their haploid genomes are inherited from the queen.

In year 1, the mean rIBD relatedness for different relatives corresponded to the mean eIBD relatedness (Table 2, Figure 2). Unlike the eIBD, rIBD relatedness takes into account recombination and segregation of chromosomes since the founders and captures distribution of realised relatedness due to these processes around the expected values (Table 2).

As shown in Figure 2, the mean IBSSingleAF relatedness for different relatives coincided with the means of eIBD and rIBD relatedness because all three metrics assumed the same founder/reference population. This calculation of IBS relatedness, IBSSingleAF, used allele frequencies from the population of founder *A.m.carnica* queens (IBSsingleAF). Despite the same mean, IBSsingleAF relatedness exhibited a larger standard deviation of values than the eIBD relatedness (Table 2). This larger variation is a result of the recent and ancestral (pre-pedigree) recombination and segregation of alleles captured by the genotype data in the IBSsingleAF calculations.

In year 10, the mean relatedness for all types of relatives increased compared to year 1 (Figure 2, Table 2), an expected consequence of a closed population. WW relationships had the largest increase in mean relatedness compared to year 1, whereas DD relationships the smallest increase, a difference which can be attributed to the honeybee’s haplo-diploidy. Workers are diploid, meaning that relatedness compares two genomes that are on average increasingly more similar over time through both the maternal and paternal side in a closed population. Notably, the numerator relationship includes a variance coefficient for each genome and twice the covariance coefficient between the two genomes Wright (1921); Henderson (1976); Grossman and Eisen (1989). Whereas, relatedness among haploid drones compares one genome exhibiting relatedness only through maternal side. We also observed an increase in the standard deviation of relatedness in year 10 compared to year 1, due to the random process of inheritance accumulating variation over time.

#### 3.2.1 Relatedness at the CSD locus within a colony

We further inspected the relatedness at the *CSD* locus. We show this in Figure 3, which we present in the same manner as Figure 2 by showing the relatedness values at the locus in years 1 and 10. The eIBD measures expected relatedness across the genome according to pedigree and therefore does not account for the balancing selection at the *CSD* locus. Hence, the eIBD relatedness at the *CSD* locus (Figure 3) was identical to the eIBD relatedness for the whole genome (Figure 2). The rIBD relatedness displayed the three possible relatedness values at the locus for worker-to-worker (WW): a relatedness value of 0.0, 0.5, and 1.0 respectively representing sharing 0, 1, and 2 alleles out of 4 alleles of two diploid workers. Namely, relatedness larger than 1.0 is not possible between workers given that homozygotes (“diploid drones”) at the locus are killed upon detection by nursing workers.

**Figure 3:**
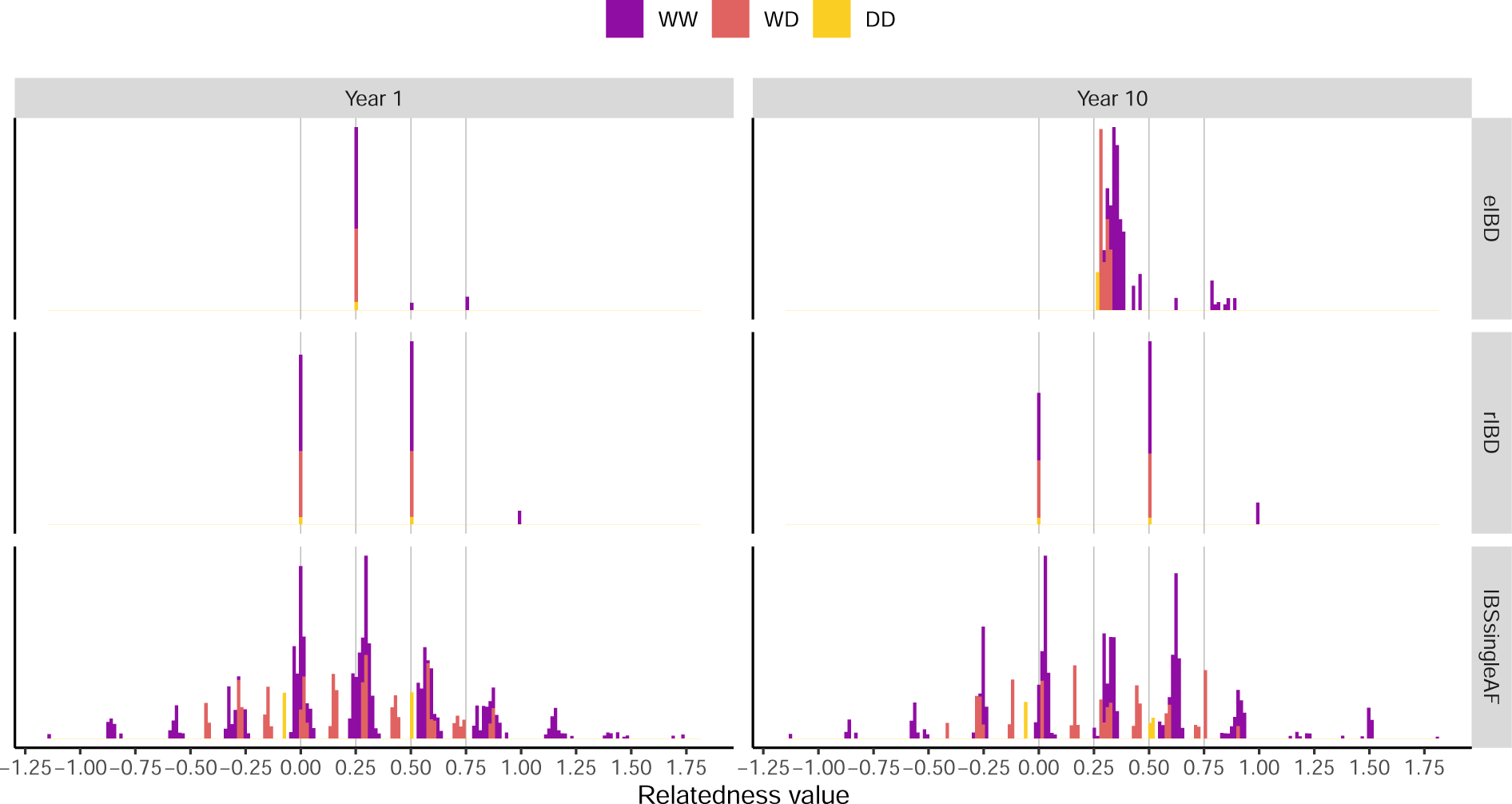
Expected and realised identity-by-descent (IBD) and identity-by-state (IBS) relatedness for the complementary sex determination (*CSD*) locus within a *A.m.carnica* colony in year 1 (left) and 10 (right) of the simulation. IBS was calculated using allele frequencies of the founder *A.m.carnica*queens (SingleAF). The figure shows relatedness between workers-to-workers (WW), workers-to-drones (WD), and drones-to-drones (DD). Vertical lines at 0.00, 0.25, 0.50, and 0.75 represent the expected within-colony relatedness of a non-inbred colony to facilitate comparisons.

In year 1, most of the workers shared 0 or 1 allele and few of them shared 2 alleles. rIBD and eIBD differed in values because eIBD reports the expected relatedness among workers from the same queen, whereas rIBD reports the realised relatedness and two workers can inherit different alleles from the same queen and hence share no IBD alleles from the maternal side. Worker-to-drones (WD) relatedness was only 0.0 or 0.5, because drones potentially share 0 or 1 allele at the locus out of 3 alleles of one worker and one drone. Note that relatedness is calculated by comparing haploid genomes’ and averaged over all possible comparisons (Grossman and Eisen, 1989; Druet and Legarra, 2020). Drone-to-drones (DD) only had relatedness of 0.0 or 0.5, because drones are haploid and therefore could potentially share 0 or 1 allele at the locus out of 2 alleles of two haploid drones (Grossman and Eisen, 1989; Druet and Legarra, 2020). IBSsingleAF relatedness had a wide distribution of values in year 1, with the mean (standard deviation) relatedness of 0.29 (0.40) for WW relationships, 0.27 (0.28) for WD relationships, and 0.28 (0.21) for DD relationship. The wide range of IBS values was a consequence of the *CSD* locus spanning 7 segregating sites, which gave rise to a large number of possible combinations of alleles at the sites and hence different IBS relatedness values.

As inbreeding increased over the next 10 years within the closed population, we observed an expected rise in allele sharing in WW and decreased frequency of WW pairs with no shared alleles, both measured by rIBD. This increase in allele sharing was also observed in WD relationships, because the queen and the fathers became more related over generations. Previous studies report that increased allele sharing, resulting from inbreeding, increases the likelihood of a lethal homozygous (spotty) brood which impacts brood numbers and possibly colony survival (Lerner, 1954; Crozier and Page, 1985; Woyke, 1980).

Because homozygous individuals at the *CSD* locus are killed, we expected lower mean eIBD relatedness compared to rIBD relatedness in year 10 for WW relationships at the whole-genome level (Figure 2, Table 2), as indicated by the mean difference between eIBD and rIBD at the *CSD* locus (Figure 3). However, we did not observe this expected difference at the whole-genome level. To explore this further, a queen was mated with her own male offspring to create a more inbred colony where progeny will be homozygous at the *CSD* 50% of time. In this case, the increased presence of homozygous diploid males resulted in a decreased mean rIBD relatedness at the whole-genome level (1.24) and at the csd chromosome level (1.19) compared to eIBD mean relatedness (1.25).

#### 3.2.2 Relatedness of the queen with her offspring within a colony

In addition to siblings, we inspected the relatedness values of the queen with her offspring (Supplementary Figure 7). In year 1, both queen-to-workers (QW) and queen-to-drones (QD) relatedness had a mean of 0.5 regardless of the metric used to compute relatedness. Both IBD values align with the paternal-offspring relationship on mammalian X chromosome described by Druet and Legarra (2020).

The relatedness of 0.5 is expected for QW relationships because workers inherit half of their diploid genome from the queen and the remaining half from their fathers (relatedness compares worker’s two genomes to queen’s two genomes), while for QD relationships drones receive all of their haploid genome from the queen (relatedness compares drone’s one genome to queen’s two genomes).

In year 10, we saw the same trend as shown in the sibling relationships (Figure 2); an increase in relatedness due to inbreeding within the closed population. Specifically, inbreeding increased similarity between the queen’s genomes and the drones she mates with, which in turn increased relatedness between the queen and her offspring. We observed a higher increase in relatedness for QW relationship compared to QD relationship, which is expected due to queens and workers having a diploid genome, drones having a haploid genome, and how the relatedness compares genome combinations of pairs of individuals.

#### 3.2.3 Comparison of IBS relatedness within a colony

We next compare IBS relatedness between different caste individuals within a purebred colony, calculated using different allele frequencies. We visualise these results in Figure 4 and present the corresponding mean and standard deviation of each relatedness calculation for different types of sisters in Table 3).

**Figure 4:**
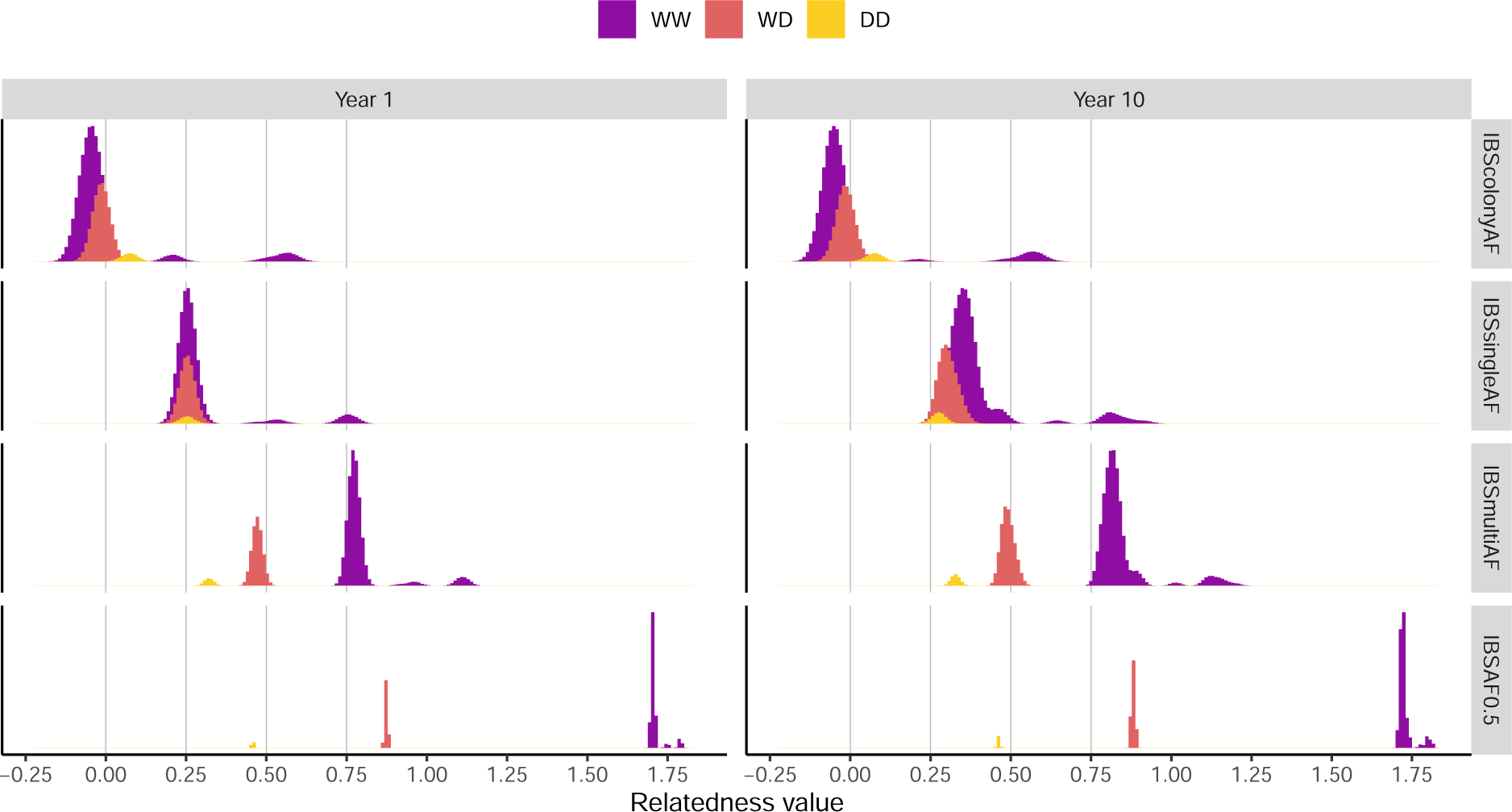
Comparison of identity-by-state (IBS) relatedness within an *A.m.carnica* colony in year 1 (left) and 10 (right) of the simulation. IBS relatedness was calculated using four different allele frequencies: the colony allele frequency (ColonyAF), the founder population allele frequency of *A.m.carnica* queens (SingleAF), the founder population allele frequency of *A.m.carnica* and *A.m.mellifera* queens (MultiBF), and allele frequency of 0.5 (0.5AF). The figure shows relationships between workers-to-workers (WW), workers-to-drones (WD), and drones-to-drones (DD). Vertical lines at 0.00, 0.25, 0.50, and 0.75 represent the expected within-colony relatedness of a non-inbred colony to facilitate comparisons.

**Table 3:** Mean and standard deviation of identity-by-state (IBS) relatedness between workers by sister type within a *A.m.carnica* colony in year 1 and year 10 of the simulation. IBS relatedness were calculated using four different allele frequencies: the colony allele frequency (ColonyAF), the founder population allele frequency of *A.m.carnica* queens (SingleAF), the founder population allele frequency of *A.m.carnica* and *A.m.mellifera* queens (MultiAF), and allele frequency of 0.5 (0.5AF).

| Year | Sister type | Mean (Standard Deviation) |  |  |  |
| --- | --- | --- | --- | --- | --- |
|  |  | IBSsingleAF | IBScolonyAF | IBSmultiAF | IBSAF0.5 |
| 1 | Half-sister | 0.25 (0.03) | -0.03 (0.03) | 0.78 (0.02) | 1.70 (0.01) |
|  | Full-sister | 0.49 (0.03) | 0.22 (0.03) | 0.94 (0.02) | 1.75 (0.01) |
|  | Super-sister | 0.75 (0.03) | 0.56 (0.04) | 1.11 (0.02) | 1.79 (0.01) |
| 10 | Half-sister | 0.70 (0.05) | -0.05 (0.05) | 1.03 (0.05) | 1.77 (0.01) |
|  | Full-sister | 0.90 (0.00) | 0.29 (0.04) | 1.17 (0.02) | 1.80 (0.00) |
|  | Super-sister | 1.04 (0.03) | 0.53(0.06) | 1.29 (0.04) | 1.83 (0.01) |

We present Figure 4 in the same manner as Figure 2, examining within-colony relatedness for WW, WD, and DD relationships in year 1 followed by year 10. We compare how IBS relatedness, using different allele frequencies, compares relative to IBSsingleAF, as described up to this point. The results show significant differences in IBS relatedness depending on the used allele frequencies.

##### ColonyAF

Alike to IBSsingleAF, year 1 relatedness between workers (WW) using the colony’s own allele frequencies (IBScolonyAF) shows three peaks (Figure 4), representing the three sister types (Table 3). However, the mean IBScolonyAF relatedness in year 1 is 0 because it is calculated relative to the colony’s own allele frequency. The mean relatedness of half-sisters indicates that they are on average less related than a random pair of individuals in the colony. In comparison, full sisters exhibit above average relatedness and even more so super-sisters. Using colony’s own allele frequencies to compute IBS relatedness shows an intriguing “flip” in drone relatedness (DD) compared to using other allele frequencies. The mean IBScolonyAF relatedness between drones is higher than the mean IBScolonyAF relatedness between half-sisters (first peak of WW), which is in contrast with other IBS calculations where drones appear less or equally related than the half-sisters. An explanation for this is that the queen serves as the common genetic denominator for the colony, representing the colony’s “internal” allele frequency. In contrast, fathers introduce “external” alleles, contributing to the genetic diversity to the colony beyond queen’s alleles. Therefore, mean IBScolonyAF relatedness among drones is higher, because they have exclusively queens’ “internal” alleles, whereas IBScolonyAF relatedness among workers (WD) and workers and drones (WD) is lower due to fathers’ “external” alleles. This difference reduces with IBSsingleAF and flips with IBSmultiAF since both use allele frequencies from founder population(s) across multiple queens, as well as with IBSAF0.5 where all allele frequencies are assumed 0.5.

It is also important to highlight that IBSColonyAF relatedness is the only method that included males in the reference population for computing allele frequencies. Since males are haploid, their inclusion leads to a reduction in the mean values of relatedness in the reference population, and an increase in the relatedness values among other individuals since they are compared to a lower average. Supplementary Figure 8 demonstrates this, where IBSColonyNM relatedness assuming allele frequency computed without considering males consistently exhibits higher mean relatedness across all relationships in comparison to IBSColonyAF.

##### MultiAF

When using the founder allele frequencies across both subspecies, *A.m.mellifera* and *A.m.carnica* (IBSMultiAF), the mean relatedness was the highest among all IBS variants. This arises because the reference/founder population we compare the colony to is on average less related compared to other reference/founder populations, such as the single subspecies population used for IBSSingleAF. Namely, for IBSMultiAF we calculated allele frequencies across two subspecies that have diverged from a common ancestor approximately 300,000 years ago (Wallberg *et al*., 2014), meaning that individual colonies deviate substantially from this common reference population giving high IBS relatedness.

##### AF0.5

Using allele frequencies of 0.5 resulted in the widest range of within-colony mean IBS relatedness (IBSAF0.5) among all the reference/founder populations. However, because the mean relatedness deviates the most from 0 we can infer that the actual allele frequency of the colony, ColonyAF, differs considerably from 0.5. IBSAF0.5 relatedness had the least variation in mean values between different sister types and the lowest standard deviation within sister type (Table 3).

##### Year 10

In year 10, the mean IBS relatedness increased for all relationships compared to year 1 (Figure 4, Table 3), regardless of the allele frequencies used in calculations, as would be expected in a closed mating population. Whilst the true increase is only one, the difference in the mean appears larger or smaller depending on the allele frequencies used for IBS calculations. The mean IBS relatedness of all variants increased the most for WW relationships, whereas DD relationships had the smallest increase as described before due to diploid and haploid comparisons. Comparing values from year 1 to year 10, we observed the largest increase in relatedness for all relatives when using allele frequencies calculated from the founder *A.m.carnica* queens (IBSsingleAF), and the smallest increase when using allele frequencies of 0.5. Using colony’s own allele frequency (IBSColonyAF) exhibited no increase in the mean relative relatedness from year 1 to 10, though the standard deviation did increase in WW relatedness (Table 3). The reason for this is that the IBScolonyAF relatedness in year 10 is based on colony allele frequency from the year 10, hence the relatedness is again relative to the average relatedness in the colony and centered on 0. On the other hand, IBSsingleAF and IBSmultiAF relatedness use the same allele frequencies in year 1 and year 10 meaning that the inspected colony is compared to the same reference/founder population.

### 3.3 Relatedness of queens within the same populations

IBS relatedness between queens within the same population is shown in Figure 5a and the queens’ to themselves in Figure 5b. IBS relatedness was calculated using allele frequencies of the founder queens containing *A.m.carnica* and *A.m.mellifera* (MultiAF). We present relatedness values from year 10 because all queens in year 1 show “background” relatedness between the most recent common ancestor and the founding population in year 1.

**Figure 5:**
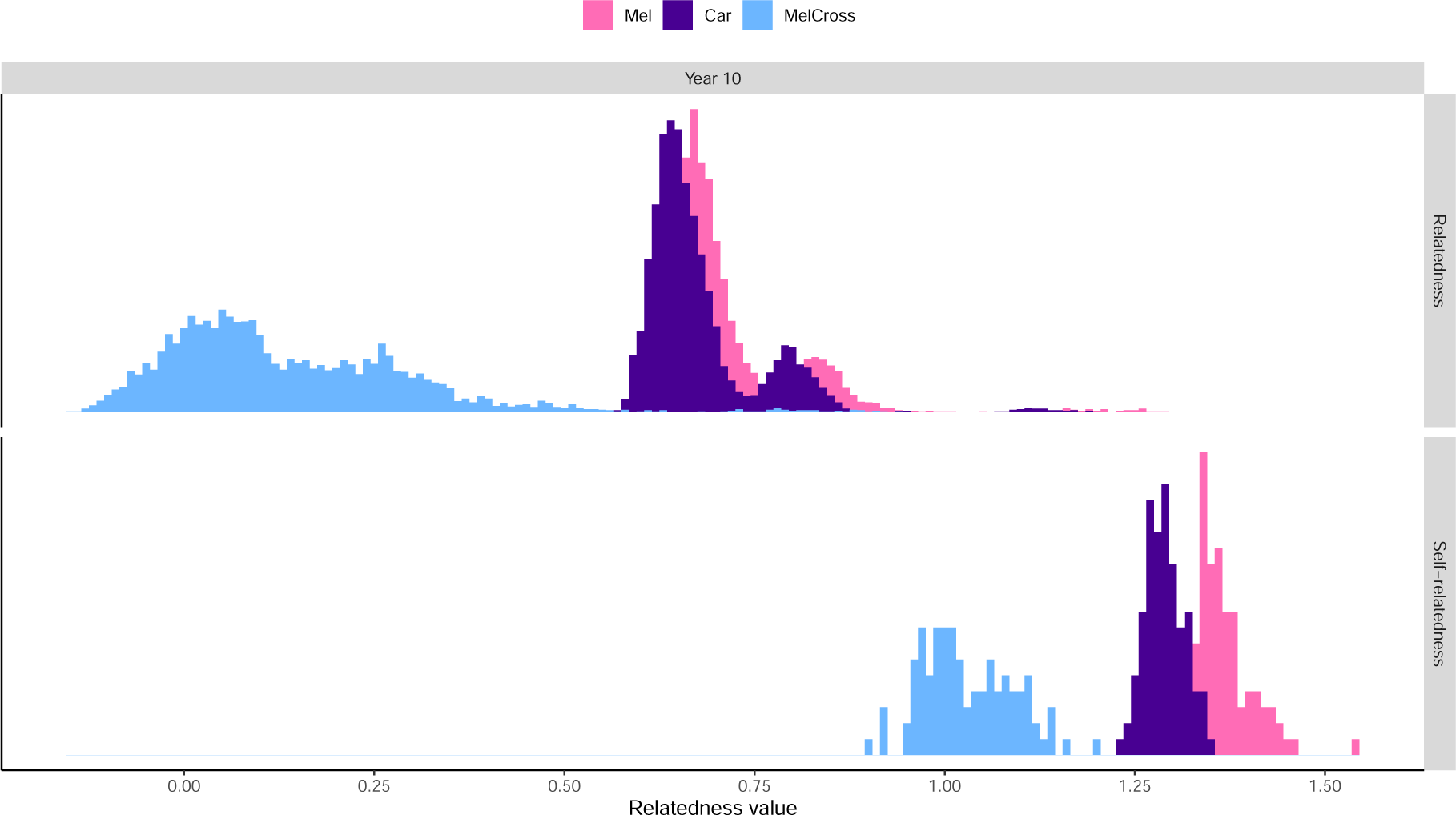
Identity-by-state (IBS) relatedness in year 10: a) between queens within two closed populations *A.m.mellifera* and *A.m.carnica* and one hybridising population *MelCross* (top-row); b) the self-relatedness of these queens (bottom-row). IBS relatedness was calculated using allele frequencies of the *A.m.carnica* and *A.m.mellifera* founder queens (MultiAF).

The hybridised MelCross queens are less related than queens in the closed populations in both between queens and in queens’ with themselves. In this simulation, we mated MelCross queens with 50% of their own and 50% imported drones from *A.m.carnica*. We used this high proportion of import for demonstration only. Given the reported UK statistics, 260,000 managed hives were accounted for in 2020 with 21,000 imported queens, hence a more realistic import percentage would be *∼* 10% (gov.uk, 2021; BeeBase, 2022). However, our results show decreased relatedness in hybrid populations compared to the closed populations. Previous studies showed that hybridisation of the populations increases the genetic variance and genetic diversity, and reduces the risk of inbreeding depression, which have all been shown to increase overall population’s health and adaptability (Brückner, 1976; Crozier and Fjerdingstad, 2001; Seeley and Tarpy, 2006).

Since both closed populations, Mel and Car, have the same number of individuals and undergo the same events throughout the simulation, we would expect the same relatedness values of both populations. In this replicate of the simulation, the Mel queens’ self-relatedness exhibit higher values in comparison to the Car queens (Figure 5b), which we concluded was stochastic, with some simulation replicates showing Car self-relatedness to be higher.

### 3.4 Relatedness of queens between different populations

The IBSMultiAF relatedness between queens of the different populations after 10 years is visualised in Figure 6 The relatedness was calculated using allele frequencies of the founder queens containing *A.m.carnica* and *A.m.mellifera*. Alike to Figure 5, we only present year 10 of the simulation in Figure 6.

**Figure 6:**
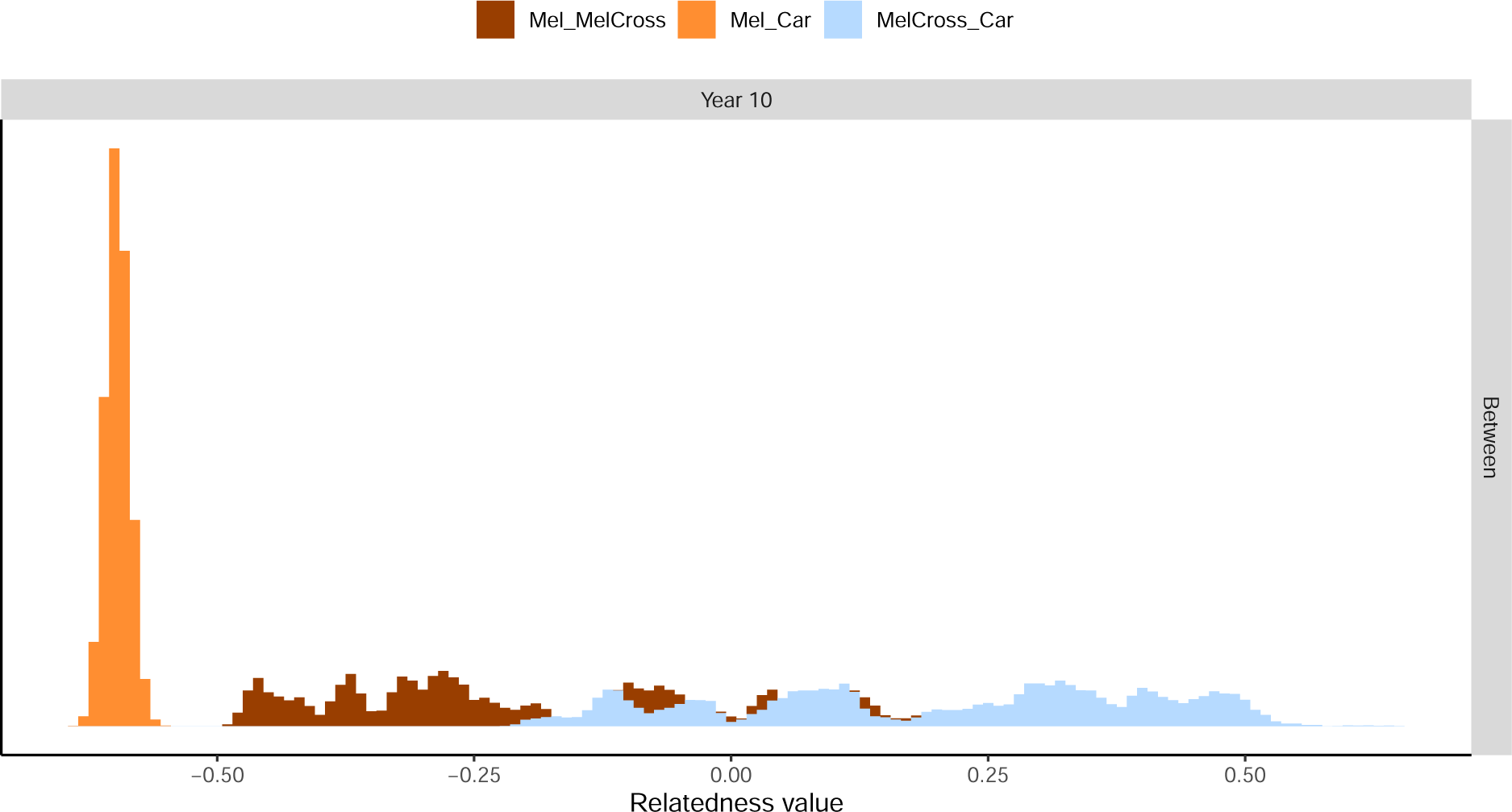
Identity-by-state (IBS) relatedness of queens between three different populations in year 10 of the simulation. Comparisons were made between the closed pop-ulations (Mel_Car) and between each closed population with the hybrid population (Mel_MelCross and MelCross_Car). IBS relatedness was calculated using allele frequencies of the *A.m.carnica* and *A.m.mellifera* founder queens (MultiAF).

We observed the lowest relatedness between the *A.m.mellifera* queens with the *A.m.carnica* (Mel_Car) queens populations (Figure 6), because these two populations have different founders and never cross. Relatedness between Mel queens and MelCross queens (Mel_MelCross) was higher compared to relatedness between Mel and Car population queens (Mel_Car). Although Mel and MelCross populations do not interact in the simulation, they come from the same founder population of *A.m.mellifera* genomes. Relatedness between queens in the hybrid MelCross population and the Car population (MelCross_Car) had the highest values. This is due to the continuous mating of MelCross queens with Car drones, introducing *A.m.carnica* through gradual accumulation within the MelCross population (Eckert *et al*., 2008).

## 4 Implications

One of the aims of this study was to investigate the impact of inbreeding and hybridisation on the relatedness of honeybee populations. Whilst reviewing the literature, we noticed a lack of studies with clear statement of relatedness metrics and underlying methods. We addressed the question both with the simulations and calculations presented in the results. These results will provide valuable insights into the genetic health and diversity of populations whilst highlighting the need for clear methods for assessing relatedness.

### 4.1 Imports debate within the beekeeping community

There is an ongoing debate within the beekeeping community around the benefits and consequences of importing queens, which impacts performance, fitness, and genetic diversity of receiving honeybee populations. Two of the most commonly imported subspecies are *A.m.ligustica* and *A.m.carnica*, given their desired gentleness and productivity in their native ranges and beyond (Lodesani and Costa, 2003). Advocates of importing queens hence argue that the lower costs of queens, gentler dispositions, higher pathogen or parasite re-sistance, and a longer period of availability outweigh any potential negative repercussions (Schiff and Sheppard, 1995; Muñoz *et al*., 2014; Bieńkowska *et al*., 2021). Imports are also argued to promote the increase of genetic diversity within the local area, however not enough studies have looked into local honeybee diversity and identified problems in the populations to claim that an increase is essential. Previous studies have indicated that such an increase in genetic diversity in honeybee populations can contribute to improved population health, honey yield, and adaptability in the fast-changing ecosystems (Tarpy, 2003; Mattila and Seeley, 2007; Simone-Finstrom *et al*., 2016). The opposing side posits that the introduction of non-native subspecies may lead to the extinction of the native population through hy-bridisation, a loss of local adaptation essential to survival in changing-environments, and to potential introduction of diseases to which native subspecies are susceptible (Moritz *et al*., 2005; Le Conte and Navajas, 2008; Byatt *et al*., 2016). Eckert *et al*. (2008) suggest that whilst hybridisation may increase local genetic variability, it could lead to an overall decrease in the global variability through reduction or even loses of local populations. Further investigations are warranted to explore the extent of this limitation in honeybees. Multiple organisations are striving to preserve and conserve native populations, maintaining their purity as much as possible. Conservation schemes focusing on *A.m.mellifera* and located throughout Europe, such as the Native Irish Honey Bee Society(NIHBS, 2012) and the Scottish Native Honey Bee Society (SNHBS, 2017).

### 4.2 Impact of the methodology

A comparative examination of different relatedness metrics revealed that the absolute re-latedness values depend heavily on the source of information used and which population is considered as a founding/reference. This underscores the importance of understanding the methodology when comparing publications and their results. While previous research has focused on other species, such as cattle (e.g. Makgahlela *et al*., 2013; Wientjes *et al*., 2017), there is a gap in the literature about this topic in honeybees. Classic genetic studies report re-latedness between individual honeybees (Brascamp *et al*., 2014; Brascamp and Bijma, 2019), while most recent quantitative genetic studies evaluate relatedness at the colony level. With the challenge of recording pedigrees in honeybees and growing use of genome-wide genotype data, we have set out to connect the classic relatedness studies with pedigrees to this new source of information. To this end, our results connect the well established eIBD relatedness based on pedigree with rIBD relatedness based on capturing recombination and segregation of genomes within a pedigree, and IBS relatedness based on genotype data. While eIBD and IBS are easily calculated from respectively pedigree and genotype data, rIBD requires identification of recombination and segregation events pedigree and genotype data, which is not trivial from observed data (e.g. Elston and Stewart, 2008; Whalen *et al*., 2018). All reported IBS relatedness metrics in this study are correct, but the assumed reference popula-tion from which allele frequencies are calculated has a large influence and care must be taken when comparing IBS relatedness metrics across studies. To calculate the IBS relatedness we used the VanRaden method 1 (VanRaden, 2008), which weighs all loci equally. There is a large number of other IBS relatedness methods (Speed and Balding, 2015), which also have implied assumptions, such as weighing loci differently according to their allele frequency (giving larger weight to loci with rare alleles) (Yang *et al*., 2010) or linkage disequilibrium between loci (equalizing weights of loci in linkage) (Speed *et al*., 2012). We have focused on individual relatedness throughout and left colony level relatedness for future work.

### 4.3 Simulation limitations

To streamline and expedite the simulation and calculation run-time, we only used 1000 segregating sites per chromosome, which likely underestimated the relatedness at the whole-genome level (e.g. Makgahlela *et al*., 2013; Eynard *et al*., 2015). Furthermore, we only created 1000 workers and 50 drones per colony. In real colonies, there can be up to 60,000 workers and a few thousand drones present in a colony at one time (Bodenheimer, 1937; Page and Peng, 2001). Whilst our numbers are smaller, they can be treated as a representative sample of a realistic setting. Moreover, we ran the simulation for only 10 years, which given the generational interval of 1-2 years is not too short a time period to see results. Our 10-year period was simply for demonstration purposes but it did enable us to show the trends of inbreeding and hybridisation. The import percentage used in this example was set unrealistically high, fixed to 50% and sourced from only one foreign population. This deliberate exaggeration aimed to highlight the impacts of hybridisation on the relatedness values within the colonies. In reality, imported animals would likely originate from numerous source populations. As described above, a more realistic import percentage would be *∼* 10% (BeeBase, 2022; gov.uk, 2021), though this number would likely fluctuate depending on the time of year and location of the hives.

## 5 Conclusions

This paper demonstrated the principles of honeybee genetic relatedness using simulation, exhibiting the potential of simulations in the study of quantitative methods as well as establishment or optimisation of honeybee breeding programmes. We evaluated the relatedness within a single colony, between queens of the same population, and between queens of different populations, demonstrating an increase in relatedness in closed breeding populations as a consequence of inbreeding as well as a decrease in relatedness within a hybrid population. We found a lot of variation in relatedness depending on the underlying method, which can lead to misinterpretations of the results and highlights the importance of understanding the methodology and assumptions made in calculations. Hence, a complete information of data sources and methodologies used is needed to avoid misinterpretations when calculating and comparing relatedness across studies.

## 6 Acknowledgements

LS and GG acknowledge funding from BBSRC ISP grants

(BBS/E/D/30002275, BBS/E/RL/230001A, and BBS/E/RL/230001C) and support from the BBSRC DTP (EASTBio) CASE PhD studentship with AbacusBio. LS, JO, JB, and GG acknowledge support from the Slovenian Research Agency’s research project L4-2624. JO ac-knowledges support from the Slovenian Research Agency’s research programme P4-0133. JB acknowledges support from the Slovenian Research Agency’s PhD studentship 1000-20-0401 and the Slovenian Research Agency’s research programme P4-0431.

## 7 Author contribution statement

GG initiated the study. GG and JO supervised the study. LS and JO developed the simula-tion, analysed the data, and drafted the manuscript. All authors have revised the manuscript and approved the final version.

## 9 Data Archiving

The simulation R package SIMplyBee is available at http://www.SIMplyBee.info with the underlying GitHub repository https://github.com/HighlanderLab/SIMplyBe and on CRAN https://cran.r-project.org/package=SIMplyBee. The simulation script is available at https://github.com/HighlanderLab/lstrachan_honeybee_relatedness

## 11 Supplementary Figures

**Figure 7:**
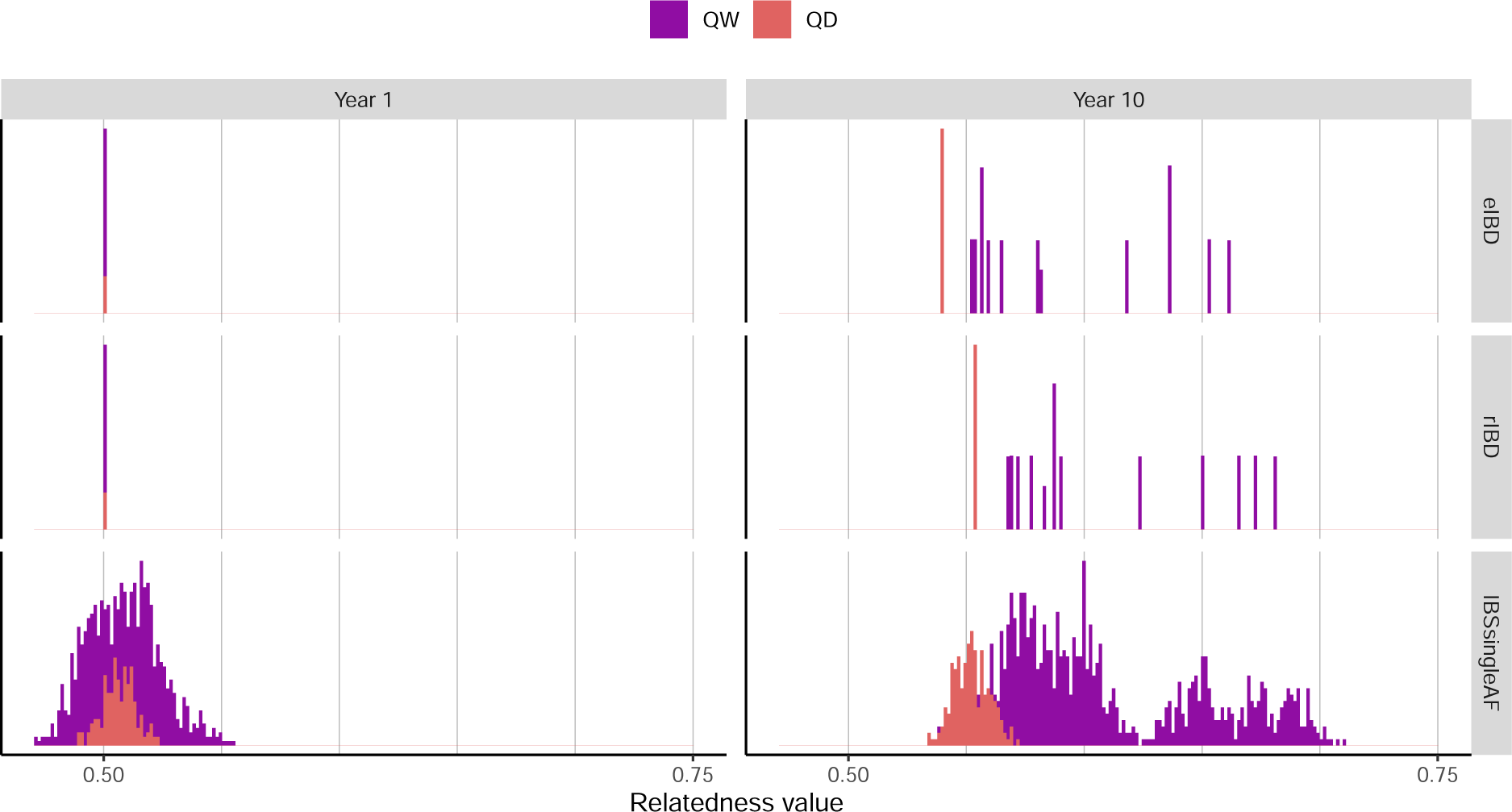
Expected and realised identity-by-descent (IBD) and identity-by-state (IBS) re-latedness within an *A.m.carnica* colony in year 1 (left) and 10 (right) of the simulation. IBS was calculated using allele frequencies of the founder *A.m.carnica* queens (SingleAF). The figure shows relatedness between queen-to-workers (QW) and queen-to-drones (QD). Vertical lines at every 0.05 between relatedness values 0.5 and 0.75 were added to facilitate comparisons.

**Figure 8:**
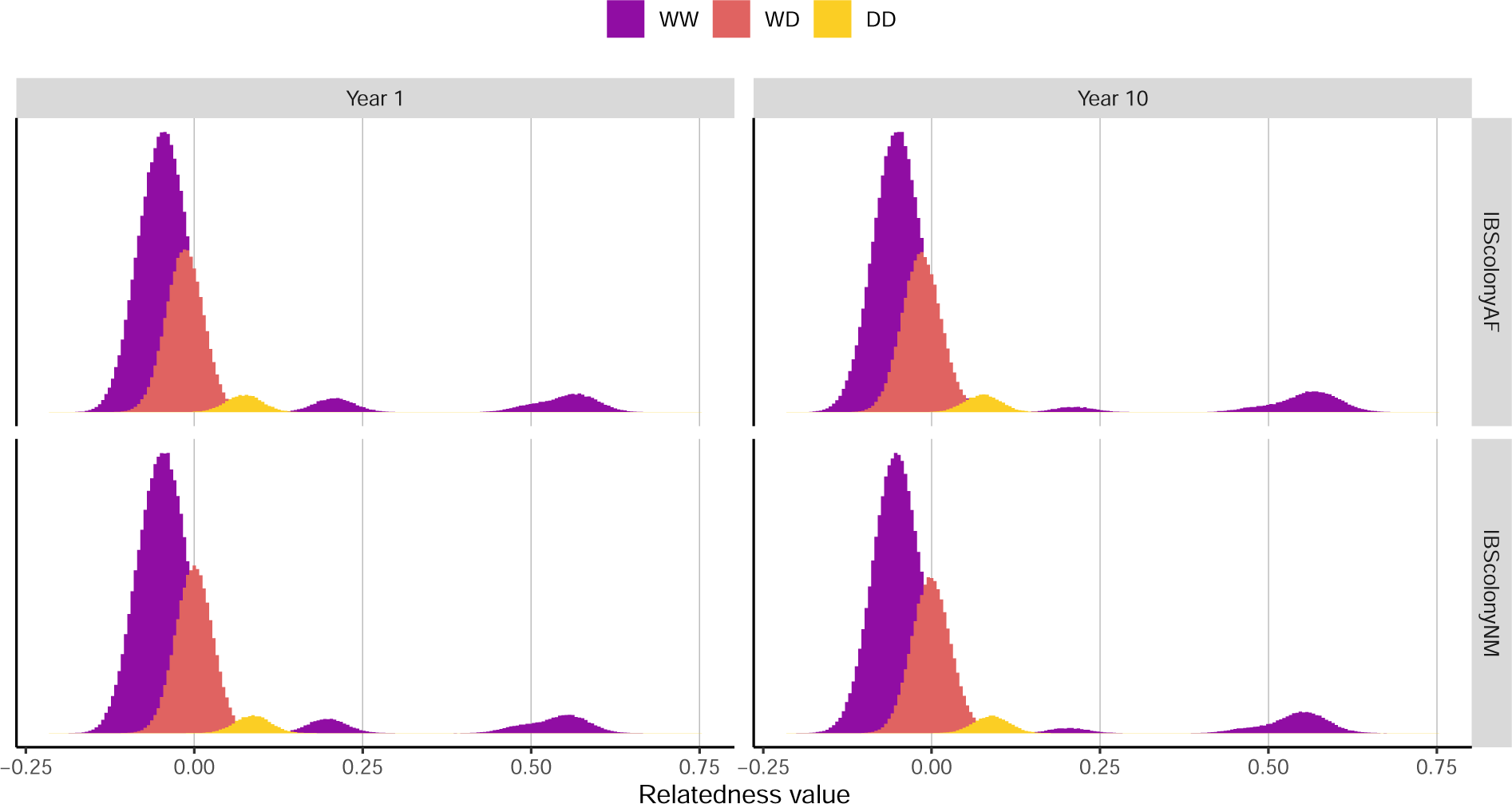
Comparison of identity-by-state (IBS) relatedness within an *A.m.carnica* colony calculated using two different allele frequencies -colony allele frequency (ColonyAF) including drones and colony allele frequency without drones (ColonyNM). The figure shows relatedness between workers-to-workers (WW), workers-to-drones (WD), and drones-to-drones (DD) in year 10 of the simulation. Vertical lines at 0.00, 0.25, 0.50, and 0.75 represent the expected within-colony relatedness of a non-inbred colony to facilitate comparisons.

